# Spatio-temporal dynamics of ingroup interactions in macaques

**DOI:** 10.1101/2025.01.10.632303

**Authors:** Tadeusz W. Kononowicz, Felipe Rolando, Lucas Maigre, Angela Sirigu, Jean-René Duhamel, Sébastien Ballesta, Sylvia Wirth

## Abstract

When sharing a space with others, many species including humans evolved a compromise regulating occupancy influenced by social determinants. For example, students in a classroom tend to sit close to their friends, keeping the same spots across days, revealing the social structure in the classroom. This place preference suggests that factors such as social hierarchy and affiliation can shape space utilization; contrasting with random walk models of agents moving at random in any given direction. Here, we asked whether spatial occupancy of macaques (*Macaca fascicularis* and *M. mulatta*) within a unisex group, reveals a structured space utilization beyond simple spatial affordance within the finite space. To this end, in two groups of four animals, we analyzed the simultaneously recorded positions of each individual while the group roamed in an enclosure. The data was gathered using automated devices that allow measuring accurate concomitant positions and calculate precise inter-individual distance, which is impossible in classical ethology even using GPS devices. Thus, our setup opens new possibilities using modelling approach, to characterize social interaction dynamics in small enclosures. We found that (1) The identity of each animal could be decoded from its individual pattern of spatial occupancy, revealing that each animal sustained a spatial footprint across multiple days. (2) Average distance between monkeys was a proxy of their social hierarchy, confirming that interpersonal distance is correlated to affiliation and dominance hierarchy. (3) Alternating the social context by removing one of the monkeys revealed that only removing the closest social partner influenced occupancy. (4) Finally, the distribution of distance between pairs of monkeys was bimodal and was modeled using random walk approach with an additional parameter reflecting propensity to stay in close proximity, which was again related to dominance hierarchy. These analyses reveal that space utilization is structured as a function of social determinants in macaques and simple modeling approach to further study group organization in neuro-ethological settings.

## Introduction

How do we manage space socially? A simple measure of the distance between two individuals can reveal their affiliation during social interaction (Hayduk 1978). Indeed, the study of inter-individual distance in humans, defined as proxemics, grounded the notion that space can be separated in intimate, personal social and public (ET Hall, 1959) and has given rise to the concept of peripersonal space. Further controlling space in their living environment, in addition to create shelters, humans often take advantage of physical borders or generate new ones, in order to create private zones which can keep the group together (houses), or divide it (rooms). Although there are cultural and socioeconomic adaptations supporting private divisions within family groups, private space remains a source of conflict within siblings in families and claim of own space is frequent within the groups (this is my seat, my room). In some respect, this parallels animal’s territorial behavior regulating space between groups in gregarious species or between individuals in solitary species (Hinsch and Komdeur 2017; Burt 1943) A behavior is considered territorial when a space is actively defended against intrusion from other animals. Territorial behavior is manifest in many non-human primates(Bates 1970). For example, it is expressed in chimpanzees known to undergo into lethal territorial encounters to defend their homeranges (Mitani, Watts, and Amsler 2010). Some other primates species, such as marmosets and lemurs are known to use chemical markings to signal their territories(Mertl-Millhollen 1988), and also enter mobbing state when two groups meet (Lazaro-Perea 2001; French et al. 1995; Caselli et al. 2018). In macaques, territorial behavior isn’t clearly manifest and neighbouring groups are known to roam through overlapping homeranges (Willems, Hellriegel, and van Schaik 2013; Bates 1970). Yet, it has been shown that territorial conflict can emerge depending on variation of resource availability seasonally (Hansen et al., 2020) or via external manipulation (José-Domínguez et al. 2015; Rebout, Desportes, and Thierry 2017). Within the group, how space is shared is actually unknow. It is known that Rhesus and fascicularis macaques show strong agonistic behavior in link to social dominance (Thierry 2007; Thierry, Singh, and Kaumann 2004) and we hypothetize that space occupancy may be a variable affected by these dominance hierarchy. Whether and how space within the group is socially distributed is less known and such information would be of importance given that these animals are the closest experimental model to human used in neuroscience studies (Feister 2018). In this study, we used the opportunity to study spatial behaviors in two groups of captive macaques housed together. Thanks to their complex social interactions that rests the use of finely grained facial expression and grooming to establish and maintain dominance (Maestripieri and Hoffman 2012; Thierry, Singh, and Kaumann 2004; Maestripieri 2007), rhesus and fascicularis macaques provide an intriguing parallel to human social behavior.

However a little is known on detailed spatial occupancy distribution within a group in macaques because although tracking of individuals with GPS has improved, positions data within a group isn’t very precise, and social determinant wasn’t often used as a variable determining space use (Xie et al. 2024; Farine et al. 2017; Chan et al. 2013). Therefore, we set out to test whether monkeys exhibited a spontaneous partition of space, and further how hierarchy and affiliation may be revealed through spatial occupancy. Since space is a limited resource in laboratory animals, we asked whether spatial occupancy can inform us about inter-individual relationships in groups of captive macaque monkeys. We determined whether spatial occupancy of macaques (*Macaca fascicularis* and *M. mulatta*) within a 2 unisex groups, reveals a structured space utilization. We used concurrently collected position data of animals in a cage to identify monkeys identity based on their spatial occupancy. We show how it is possible to access the dynamics of individuals in a group. Animals form inner spatial preferences within the cage enclosure showing that space is socially distributed and further the dynamics of interactions also reveals affiliative pattern within a group.

## Results

### Monkey identity can be decoded from its spatial occupancy

Via video-based analysis, we computed the simultaneous positions of four female macaques (Group A) co-housed together (Ballesta et al. 2014, see methods), while they were roaming in an enclosure equipped with video cameras (**Fig. 1A** and **Fig. 1B**). Data from a second group of 4 males (Group B) were also collected in the same enclosure independently (Supp Fig 1 A). In both groups, the animal’s individual status in the dominance hierarchy (1st to 4rth) was estimated during competitive encounters in parallel to the recording sessions (see methods). Averaging movement trajectories for each monkey readily revealed pattern of spatial organization (**Fig. 1CD, Supp.** Fig. 1BC) whereby separation between average positions for each monkey is notable. Visualization of data using low dimensional manifold embedding techniques suggested the same conclusion as the data points representing 1^st^ and 2^nd^ monkey where clearly grouped near each other (**Fig. 1C; Supp Fig.1C**) and away from data points representing 3^rd^ and 4^th^ monkeys, which tended to be in vicinity of each other. These two ways of visual inspection of data suggest association between space occupancy and monkey identity.

**Figure 1.**
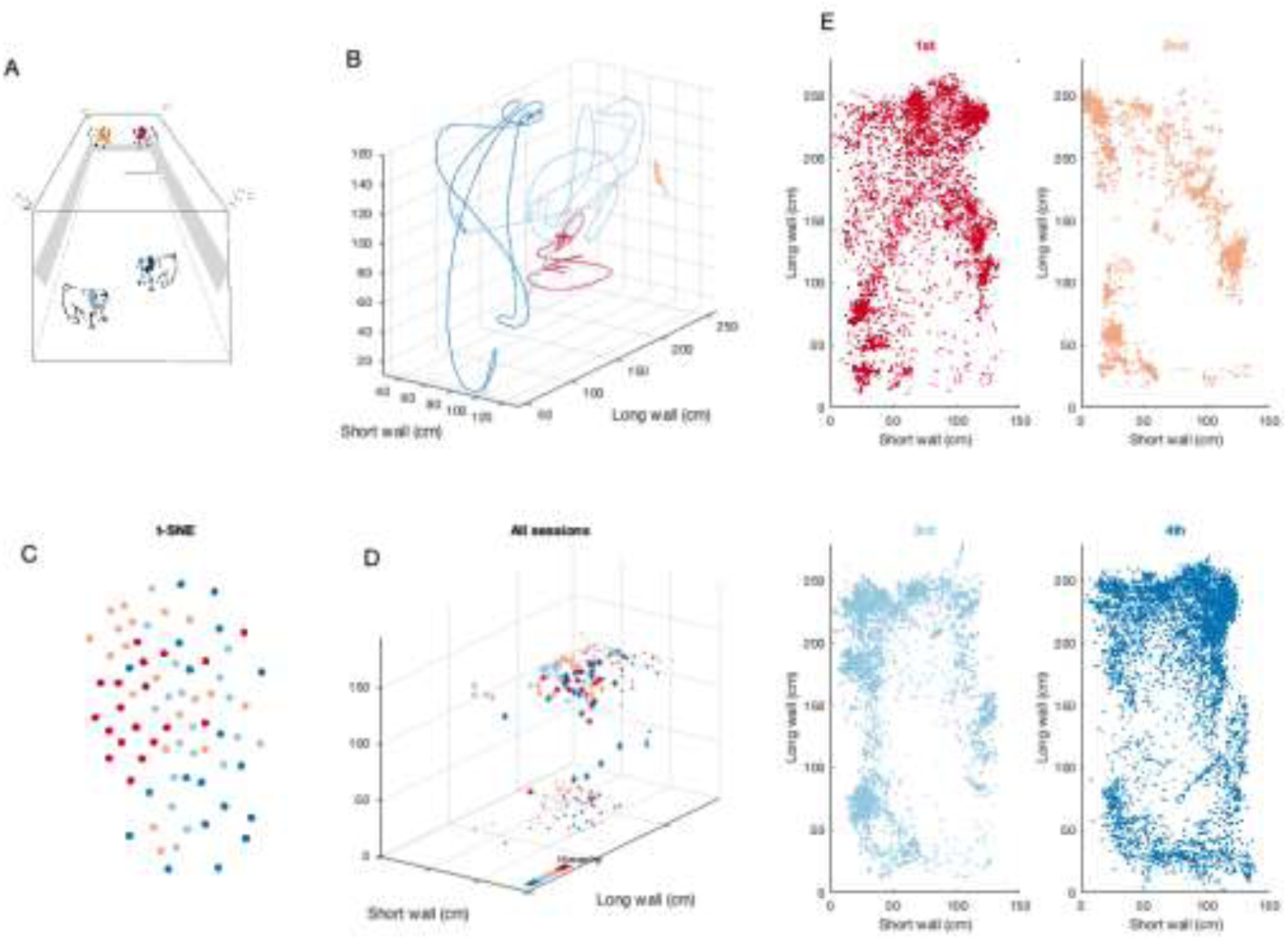
Spontaneous territorial partition of space in a group of four female macaques. (A) The home cage and tracking system design. (B) The view of the home cage. Example trajectories in 3D of four monkeys from 3 minutes of recording. The data were smoothed with 10^th^ order Savitzky-Golay smoothing filter. (C) To visual the spatial occupancy data across animals and recording sessions the data was projected onto 2 dimensional manifold using tSNE method. Single data point represents data of one individual in a single session. The main observation is that there are not scattered randomly but rather clustered. (D) Median positions per session (3h) per monkey plotted in 3D. The smaller data points depict data projection on the respective wall. Monkeys are coloured according to their hierarchy, from top to bottom, red to blue, respectively. The median positions already appeared slightly clustered, suggesting the presenceof structured distancing of monkeys in line with their hierarchy. (E) Example data from one 3 hours sessions for four monkeys recorded simultaneously. The visual inspection of patterns suggest that it should be possible to decode monkey identity based on its spatial position.

In addition, we assessed averaged distances between pairs of monkeys over sessions using ANOVA comparisons across each of three dimensions (**Fig. 1D**). A Friedman’s test showed that there was significant difference in average monkey’s occupancy across sessions in short horizontal, and vertical wall dimensions, respectively: χ^2^F(3, 56) = 18.0, *p* < 0.001; χ^2^F(3, 56) = 30.1, *p* < 0.001, besides long horizontal wall χ^2^F(3, 56) = 5.7, *p* = 0.13. This result suggest that monkey’s occupancy is not random. We hypothesized that dominance hierarchy may be one of the factors that influences monkeys’ pattern of occupancy.

In line with previous analyses, visualization of data from a single example session showed different data patterns across individuals. To quantify the informativeness of this high resolution spatio-temporal patterns, we asked whether the identity of each animal could be decoded from its individual occupancy pattern within recording sessions. To this end, we trained the Support Vector Machine (SVM, Fig. 2A) classifier on the simultaneously recorded monkey positions using 3-dimensional data recorder throughout a session. We performed three kinds of decoding analyses. First, using quadruple sessions we trained and tested classifiers within each of quadruple sessions inside 10-fold cross validation (**Fig. 2BC; Supp.** Fig. 2A**- C**). Significant decoding scores signify the fact that occupancy patterns generalize within a single session (**Fig. 2D, Supp.** Fig. 2B, 0.82-0.85 accuracy for all pairs). Second, to access whether the occupancy patterns generalize across days we trained a decoder on the data from entire session and tested on the all-remaining sessions (**Fig. 2E, Supp.** Fig. 2C). Decoding accuracy was above the chance level for all monkey pairs (**Fig. 2E**, right panel, **Supp.** Fig. 2C**, right panel**), showing that each animal sustained its spatial footprint across multiple days. Note that results were qualitatively similar in the second group of macaques (**Supp.** Fig. 2).

**Figure 2.**
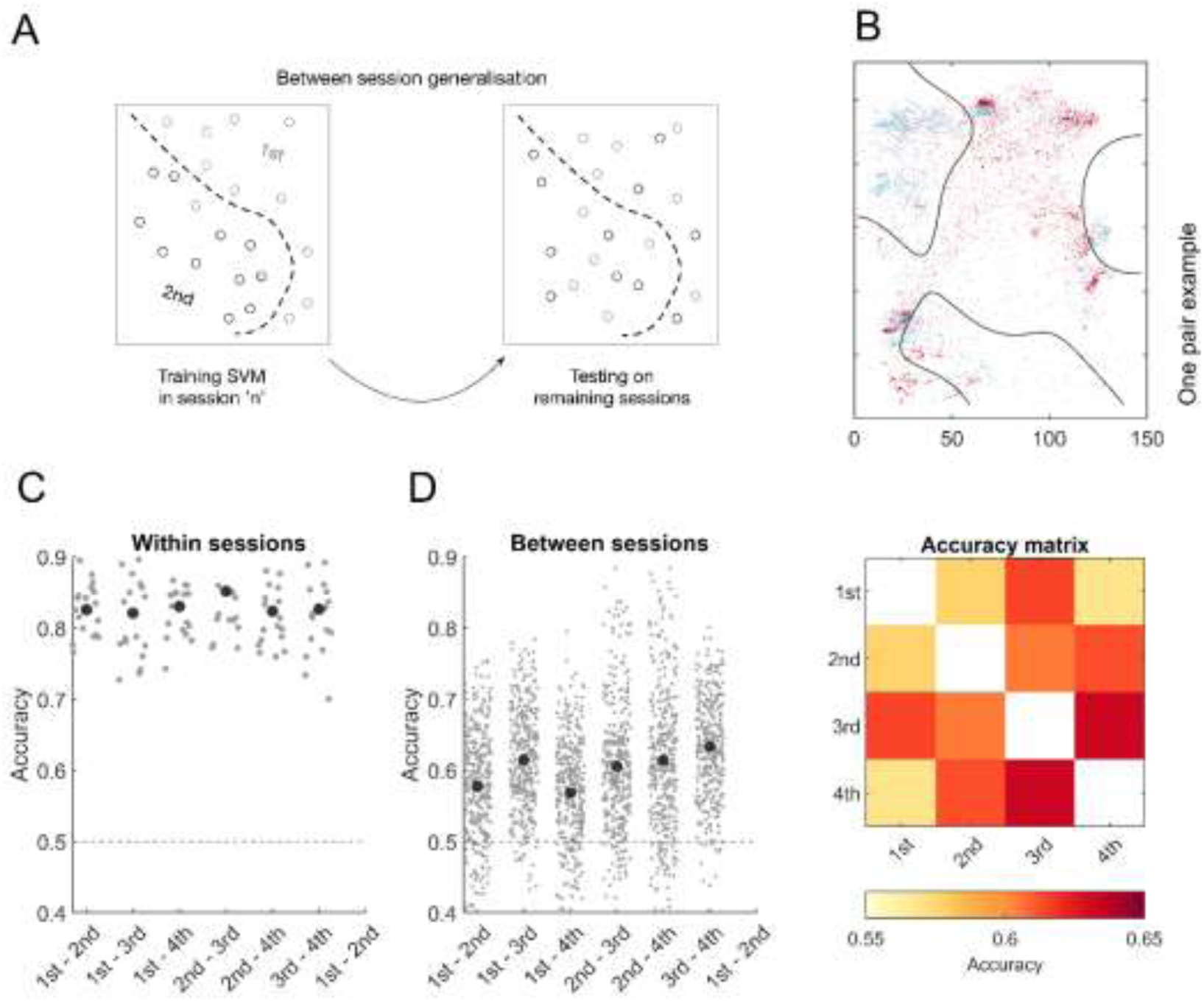
Decoding monkeys identity based on spatial occupancy reveals structured pattern of spatial utilization. (A) Schematic of decoding pipeline of monkey’s identity from their spatial occupancy pattern. We trained the Support Vector Machine classifier (SVM) on the simultaneously recorded monkey positions using 3 dimensions. Using simultaneous position across time recorded for each monkey pair we decoded monkeys identity such that each classifier was trained to distinguish between a pair of monkeys. (B) Example of one such trained classifier. The trained classifier was tested in a cross validated manner (panel D) or on all remaining 18 recording sessions (panel E). (C) To decode spatial occupancy within each session and monkey pair cross-validation procedure was performed. Decoding accuracy was above the chance level for all monkey pairs (bottom left panel), showing that each animal sustained its spatial footprint within each session. (D) The same decoding was performed on data across sessions which showed similar pattern of results. Decoding accuracy was above the chance level for all monkey pairs, demonstrating that each animal sustained its spatial footprint across multiple days. Average accuracy scores for each monkey pair are summarized in matrix form in the bottom right panel.

Next, for the group A, we investigated the influence of social context by comparing monkey’s positions in the absence of a cage-mate on different sessions (Triplet sessions) relative to sessions in which all 4 animals were present (quadruple sessions). This created different social context for monkey under consideration by alternating her neighbors. We assessed how the classifier trained on a given pair of monkeys in quadruple context performs on the same pair of monkeys in the triplet context (**Supp.** Fig. 3). Decoding accuracy higher than chance level showed that each animal sustained its spatial footprint across multiple days, even when the context changed.

### Quantifying social influence from spatial occupancy data

We aimed to further characterize how the absence of each individual impacts the spatial occupancy of other individuals during the triplet sessions (**Fig. 3**). One possibility is that only neighboring monkeys in term of hierarchy would be influenced by the absence of a given monkey. The alternative is that all monkeys would be affected by the absence of a given individual.

**Figure 3.**
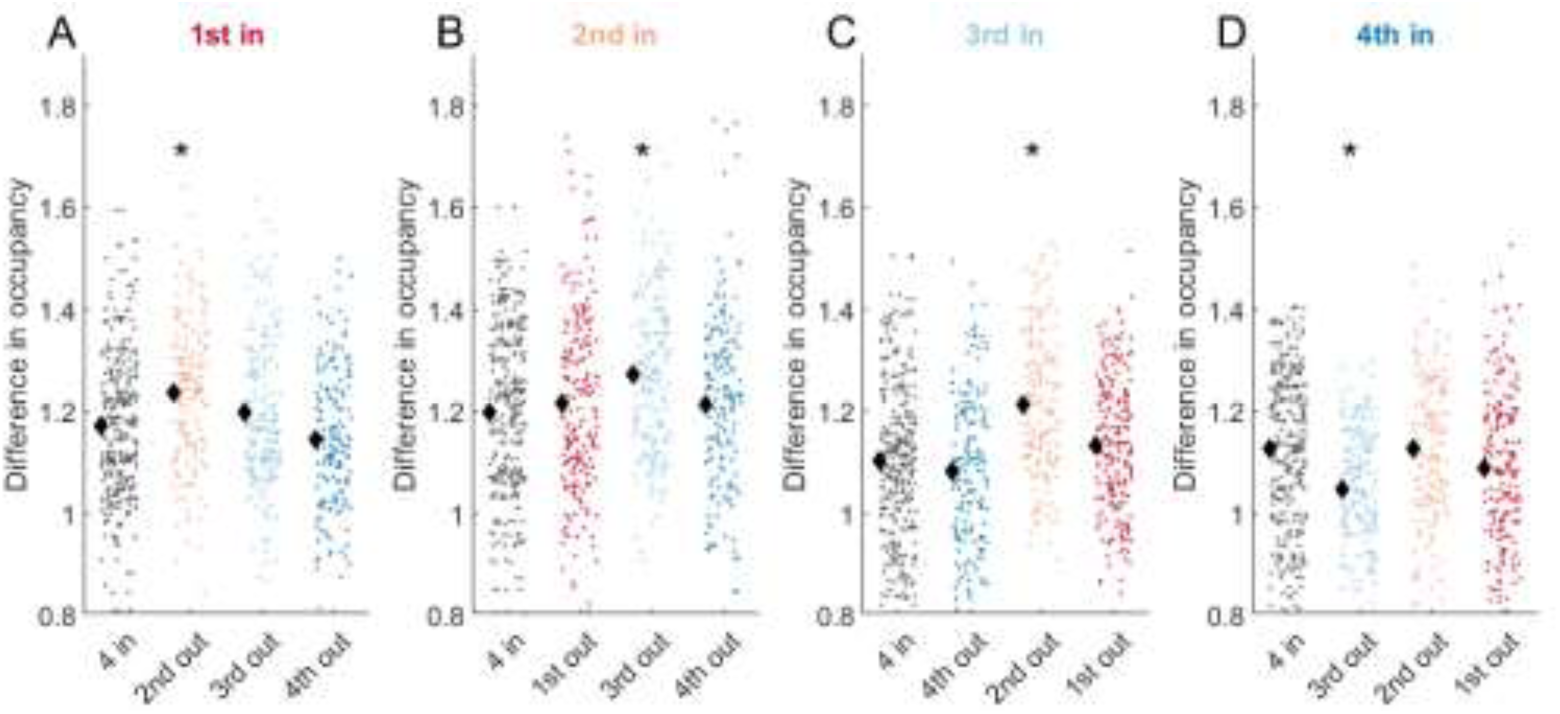
Social context influences monkeys occupancy. Every panel displays the difference in occupancy for sessions with all for monkeys in the home cage (’4in’), and three kinds of triplets where one of the other three monkeys was removed, therefore creating an effect of social context for monkey under consideration in one of the four panels. Single data point has been computed by absolute difference between normalized occupancies from two different sessions originating from the same subject, according to the following formula: mu(|NCn-NCm|), where NC is a 16 by 29 matrix of time spent in a given location normalized by total duration of a recording session.

Every panel of **Fig. 3** represents data of a particular monkey under different experimental conditions. To summarize the change between the quadruple (‘4 in’) session and triplet session in a single number, we computed L1 normed difference between normalized occupancy maps, which we denote as *D*. Essentially, a single panel in **Fig. 3** depicts the amount of change in a given condition with respect to quadruple sessions; except the first column of each panel of **Fig. 3A**, which captures to what extend the quadruple sessions vary with respect to other quadruple sessions for a given individual. The latter serves as a baseline level for comparing other conditions. To evaluate the impact of each individual on other’s spatial occupancy, we compared values of *D* between ‘4 in’ sessions and the remaining sessions (colored columns). Significantly different pairs are indicated by asterisk mark. The 1^st^ monkey in the hierarchy significantly changed its occupancy when 2^nd^ monkey in the hierarchy was absent (*p* < 0.02; **Fig. 3A**). However, 2^nd^ monkey did not significantly change its occupancy when 1^st^ was absent (*p* > 0.1; **Fig. 3B)**, but instead, was affected by the 3^rd^ monkey’s absence (**Fig. 3B**, *p* < 0.01). Symmetrically, 3^rd^ monkey was affected by the 2^nd^ monkey’s absence (**Fig. 3C**, *p* < 0.001). Finally, 4^th^ monkey was only affected by the 3^rd^ monkey’s absence (**Fig. 3D**, *p* < 0.01, fourth panel).

Overall, social context manipulation allowed us to show that monkeys were modestly, yet significantly influenced only by their closest neighbors’ absence, while other monkeys kept steady occupancy structure. Depending on which neighbor (higher or lower in hierarchy) exerts its influence gives a tool to assess symmetry and directionality of that influence.

### Time spent together as a proxy of affiliation

While spatial occupancy of co-housed monkeys revealed individual spatial patterns, we hypothesized that that the interpersonal distance between monkeys may reveal dominance and affiliation structure of the group. First, we computed the overall average distance between monkeys for each session (**Fig. 4A**, left panel**; Supp.** Fig. 4A) with a single entry of the matrix showing average distance between a pair of monkeys. Using t-tests (see methods), we compared the series of instantaneous distances between monkeys for different pairs. All combinations of pairs significantly differed besides a difference between 1^st^-4^th^ and 2^nd^-4^th^ (t(18) = 1.9, p > 0.1), suggesting that the 4^th^ monkey was equally distant from the first two monkeys in the hierarchy. To illustrate the physical distance between monkeys as a function of the hierarchy, we rearranged distance matrix in the right panel of **Fig. 4A** (right panel). In this group, averaged distance increased as the dominance hierarchy distance increased as well. The analyses of the second group of macaques revealed qualitatively the same results (***see Supp Fig 5***).

**Figure 4.**
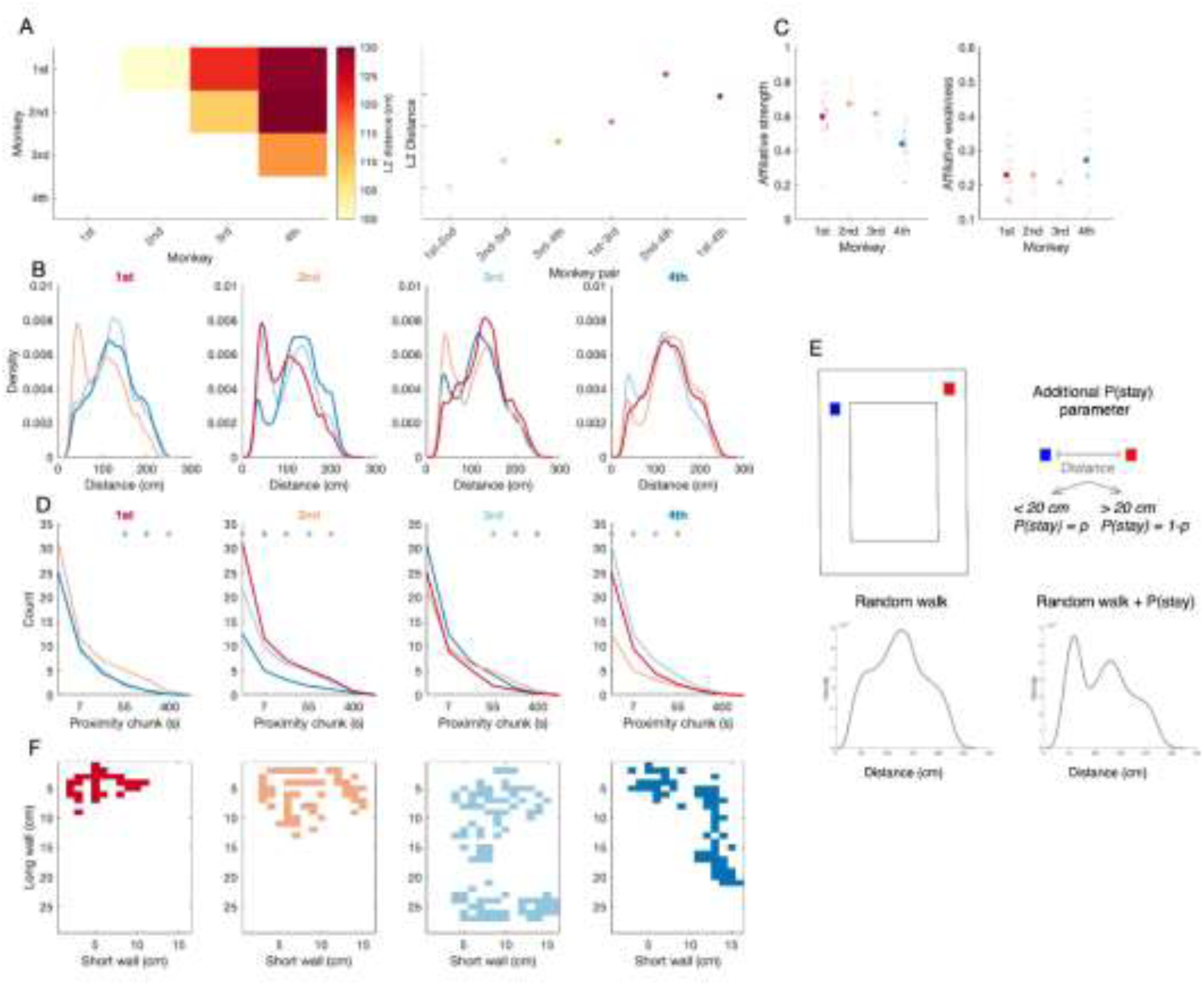
Features of interpersonal distance as a proxy of social structure. (A) The matrix represents the average distance for all sessions (n=19) for each pair of monkeys depicted in the matrix form. The redness of each cell represents average distance between a pair of monkeys. Higher values are obtained for 1^st^ and 4^th^ in the hierarchy (top right). The right panel depicts that the physical distance increases as the rank distance between any two monkeys increases. (B) We computed Euclidian distance between each pair of monkeys over time. Individual density distribution depicts distribution of distance between given monkey pair. For example, the first panel displays three density functions which depict distribution of distance between 1st monkey and other three monkeys. The same follows for the three other plots. (C) Affiliative strength and affiliative weakness calculated as total time each monkey spent with all other monkeys or total time spend away from other monkeys, respectively. (D) Temporal distribution of close proximity chunks. The same color coding follows as in the panel A. The top stars indicate the samples where significant differences have been found. (E) Schematic and results of random walk model capturing bimodal distributions. The top panel shows schematic of simulation of two agents. The bottom panels display example simulation involving only random walk and the second one involving random walk with additional social rule of behavior. (F) Map of locations where monkeys was separated from the other monkeys by at least 120 cm.

Next, we examined the distribution of interpersonal distance for each pair of monkeys at a fine resolution (**Fig. 4B** for the group A, and **Suppl Fig 5** for group B). For each group, there are two peaks in the distribution of distances for at least one pair of interactions per monkey, with the left peak expressing the amount of time spend in close proximity within the given pair. Indeed, significant differences were localized in the left part of interpersonal distribution (**Supp.** Fig. 5**, Supp Fig 6**). In the group of females, the monkey at the top of hierarchy spent a lot of time with the next monkey in the hierarchy. Strikingly, the monkey with the lowest rank spent very little time with other monkeys. In the second group, an overall similar pattern was observed with the presence of a left sided peak for at least one pair of interactions. Contrary to the group of females peak of close proximity has been developed between 1^st^ and 3^rd^ monkeys (**Supp.** Fig. 4B). In addition, some males (3^rd^ and 4^th^ monkeys) unexpectedly showed large right sided peak suggesting avoidance.

The time spent in close proximity with others, can vary from one monkey to another depending on affiliation. To capture differences between animals; we calculated the time spent in close proximity with others (i.e. less than 80cm away, defined as affiliative strength (**Fig. 4BC**, left panel). In the females, the affiliative strength of 4^th^ monkey was smaller than that of other monkeys (1^st^: Z = 3.74, *p* < 0.001; 2^nd^: Z = 4.03, *p* < 0.001; 3^rd^: Z = 3.88, *p* < 0.001). The 2^nd^ monkey tended to have the strongest affiliative strength, which was larger than affiliative strength of 4th monkey (Z = 4.03, *p* < 0.001) and nearly 1^st^ monkey (Z = 1.69, *p* = 0.09).

Conversely, the analysis of affiliative weakness, that is the time spent away from others (i.e. above> 180 cm), revealed that the 4^th^ monkey spent the least time with the other monkeys (1^st^: Z = 1.66, *p* < 0.096; 2^nd^: Z = 1.84, *p* = 0.066; 3^rd^: Z = 2.19, *p* < 0.05).

Interestingly, these results corroborate the findings of social context manipulation where 2^nd^ monkey had the most influence over other individuals (**Fig. 3**). The second observation in line with the findings from social context manipulation is that the proximity peaks occur for the monkeys further away in the hierarchy. For the monkeys further away in the hierarchy the close proximity peak is almost negligible. In sum, analysis of affiliative strength adds another dimension to analysis of dominance hierarchy. A similar pattern regarding affiliative strength and weakness was corroborated by the analysis of the second group of male macaques (**Supp.** Fig. 5BC)

The analysis above captures affiliation via the computation of overall time spent together, but does not capture the temporal dynamics of affiliative behavior: when monkeys are close together, how long does this last? To address this question, we analyzed the duration of the bouts of time (chunks) spent in close proximity and analyzed chunk size distribution (**Fig. 4C, Supp.** Fig. 4C). It is easy to notice that the chunk size distribution follows the scaling, such that for all pairs of individuals, the number of short interactions is higher than the number of long interactions. Interestingly, different pairs differed in the frequency of those short and long

interactions (**Fig. 4C, Supp.** Fig. 4C). From this analysis it becomes evident that the peak of close proximity is driven by the large number of long interactions as opposed to large number of short interactions in both groups of monkeys.

### Accounting for distribution of interpersonal distance

To capture what accounts for the nature of interpersonal distance distribution (**Fig. 4A**), we took a mechanistic approach, by first testing whether the right side of the distribution could be simulated as free roaming behavior. This can be modeled as a two-dimensional random walk process (Codling, Plank, and Benhamou 2008; Bartumeus et al. 2005; Bovet and Benhamou 1988); *Methods: Random walk model simulation*). While the random walk model captured well the right side of the distribution of interpersonal distance (**Fig. 4E**), it did not capture the peak of close proximity for some monkey pairs. Further, previous analyses suggested that occupancy was not due to purely random walks. Therefore, we included an additional parameter in the previous simulation, expressing the propensity to stay in close proximity once two agents were in close proximity. Precisely, when in interpersonal distance was below the threshold, next positions were kept for the next sample according to the parameter’s fitted probability (see methods). This simple simulation captured well the distribution of interpersonal distances for pairs showing a bimodality in their distribution of pairs (Figure supp 4). The strength of this parameter was related to distance in dominance hierarchy in the females (**Fig. 4E**, bottom panel), yet this was less the case in Group B.

### Distribution of ‘private’ spots

The analysis of interpersonal distribution showed differences in how monkeys stay in proximity or far from each other in different pairs. To tie back these results to spatial dimension, for each monkey, we identified the places where monkeys stayed away from all other monkeys (**Fig. 4E**). The maps show spatial distribution of individual space where given monkeys tend to stay on their own. Overall, individual spaces for individuals were non overlapping with provides more evidence for structured space utilization in macaques. In both groups, spots in which the 1^st^ and 2^nd^ animals in the hierarchy stay alone, tended to overlap.

## Discussion

Capturing natural social interactions necessitates by essence an understanding of live interactions. In any non-human primate society, these interactions are going to take place in a real physical space, and individual’s positions and distances with respect to each other may reveal the nature of interaction (Farine et al. 2017). Here, we took the opportunity to analyze a very simple measures in two groups (rhesus and fascicularis: by analyzing the concurrent spatial positions of individuals, we asked how space is jointly shared within the groups. We showed that there is a spontaneous distribution of space between individuals such that animal’s identity could be decoded from their position (Fig. 2D). While animals could have displayed a homogenous occupancy, we showed this was not the case. Further, individual’s spatial preferences remained sufficiently stable to decode individual’s identities across different days (Fig 2E), and decreased only when decoding was applied on sessions during which closest monkey in the hierarchy was absent (Fig3). Next, we showed that all monkeys displayed social preferences expressed as an increased time spent to some other animals over others (Fig 4). The close proximity for pairs with a strong affiliation, consists of a few relatively long-time bouts (∼6minutes) during which animals stay close, while most of the time, monkeys spent less than a second in close proximity. The bimodal distribution of interpersonal distances expressing these individual social preferences could be modeled by simple parameters which is the probability to stay together, added to the simulation of a random walk afforded by the physical structure. While network analysis is usually based on the frequency of interaction, here we show that simple measures of interpersonal distance reveal the complexity of the social network intertwining dominance hierarchy and affiliation.

## A spatial footprint for individuals

Macaques aren’t known for a strong territorial behavior, and neighboring groups tolerate overlapping home ranges(Thierry, Singh, and Kaumann 2004; Hansen et al. 2020). Further, increase in group size did not impact interpersonal distance within the group (Xie et al.,2023), but increased daily path length, suggesting the latter is a coping strategy relative to heightened competition in larger groups. Here, we we reveal that in absence of food restriction, a simple metric such as individual’s position within a group, within and across days, supports decoding individual’s identity. It is also on par with some previous work showing that animals have preferences for specific locations within a space(MacLean et al. 2009) but our data shows that animals express individual preferences akin to a signature footprint. We have some evidence suggesting that individual spatial occupancy may be reveal hierarchy, as i) decoding an individual’s identity was most affected when the closest animal in the hierarchy was absent, and ii) locations in which the 1^st^ and 2^nd^ animals in the hierarchy were “alone” were similar across the two groups of macaques. First, this suggest that the spatial occupancy of a monkey changed the most when its closest partner was absent. This suggest some sort of pyramidal control in which behavior is affected by close rather than distal interactions. Second, the preferred position physically consisted of an ideal observation spot in the cage from which the whole room could be assessed visually. Therefore, the results suggest that dominance hierarchy, in addition to allow access to food resources, also regulates access to spatial zone of interest. .

## Interpersonal distance reveals affiliation

In non-human primates, allogrooming is central to maintaining social bonds (Dunbar 1991; McCowan, Beisner, and Hannibal 2018) and reveals the structure of the social network (Seyfarth 1980; Matheson and Bernstein 2000; Simons et al. 2022). Here, we show that interpersonal distances reveal affiliation without the cost of scoring grooming as is usually performed. Specifically, the distribution of the interpersonal distance is bimodal for some pairs compared to others, showing that 1) there is a “one arm length” distance that emerges naturally from the distribution, and 2) animals had individual preferences to stay close to some animals over others. The bimodal distribution could be modeled by adding a simple parameter to a random walk, such that the probability to remain close was increased when interpersonal distance reached a specific threshold. This shows that the bimodal distribution is not the results of distribution of movements by physical affordance, but reveals specific affiliations. These specificities were present in both groups of males and females, although in the male, there were also tri-modal distribution, suggesting that some males also avoided other males and tried to stay the further away from other in the enclosure. This is in line with results showing that females invest more time in allogrooming than males as a mean to alter or support dominance hierarchy (Seyfarth 1977; Schino 2001). The analysis of the durations spent together revealed that for each pair, there were a few bouts of time during which animals stayed together. Specifically, to our surprise, animals rarely spent more than 6 continuous minutes in close proximity. The times during which the data was collected was in the late afternoon while the staff had left and animals were not disturbed. So, our data should reflect monkey’s naturalistic behaviors. While our data is consistent with previous findings showing that females gave or received grooming for about 15 minutes per hour (Martel et al. 1995; Schino 2001). We show that the time spent in close proximity to others, which may include grooming moments, is highly dynamic even in captive monkeys, and is interspersed with other activities.

## Limitations of study

A limitation of this study is the small sample size and focus on only two macaque groups, which limits broader comparisons. However, performing experimentation on social context in larger groups would be extremely difficult. GPS data may not provide the level of resolution needed to get access on fine interpersonal distance that is provided in this study. Our study is only based on two groups. Despite that they differed by sex, age and species, we found consistancies between in results. However, replication across many groups may allow to generalize the findings beyond our small sample. However, the study provides valuable insights into sociospatial behavior, and future research with larger samples and more groups could further enrich our understanding of these dynamics with the proposed tools.

## Conclusion

Overall, the study shows that the dynamics of spatial occupancy reveals the nature of social interaction in macaques consistent with social hierarchy and affiliation. One of the strength and originality of the paper that can be put forward is that usually territory or range are studied at the group level while here this is studied at the individual and dyadic level. As our study was robust across groups and across sessions, our method provides a simple and accessible tool to objectify social relationships within a group revealing interesting interindividual patterns that structure the social network.

## Acknowledgements

This work was supported by ERC socialeyes (JRD), ERC oxitocinspace attributed (AS and SW), ANR-11-LABX-0042/ ANR-11-IDEX-0007 to SW, JRD and AS.

---

**Supplementary Figure 1.**
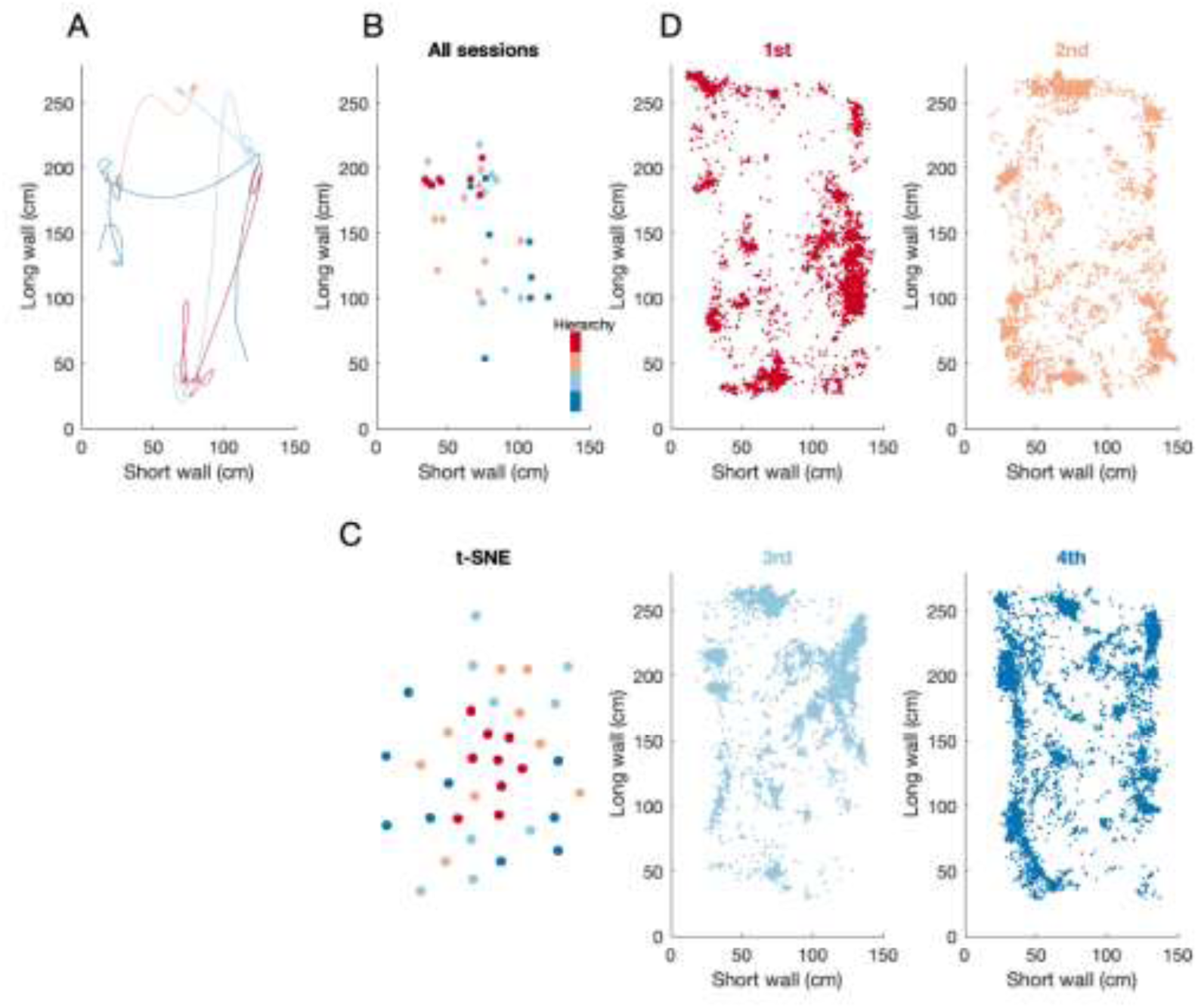
Spontaneous territorial partition of space in a second group of four male macaques. (A) The floor view of the home cage. Example trajectories of four male monkeys from 3 minutes of recording. The data were smoothed with 10^th^ order Savitzky-Golay smoothing filter. (B) Median positions per session per monkey plotted in a floor view. Monkeys are coloured according to their hierarchy, from top to bottom, red to blue, respectively. Clustering of median positions suggested that we should have observed structured distancing of monkeys in line with their hierarchy. As for the group of females, we assessed averaged distances between pairs of monkeys over sessions using ANOVA comparisons across each of three dimensions. A Friedman’s test showed that there was significant difference in average monkey’s occupancy across sessions in short wall dimensions: χ^2^(3, 24) = 11.9, *p* = 0.008; but not for long and vertical dimensions, respectively: χ^2^(3, 24) = 6.2, *p* = 0.102; χ^2^(3, 24) = 5.4, *p* = 0.144. (C) tSNE decomposition in a group of male macaques. Single data point represents data of one individual in a single session. The main observation is that there are not scattered randomly but rather clustered. (D) Example data from one sessions for four monkeys recorded simultaneously. The visual inspection of patters suggest that it should be possible to decode monkey identity based on its spatial position.

**Supplementary Figure 2.**
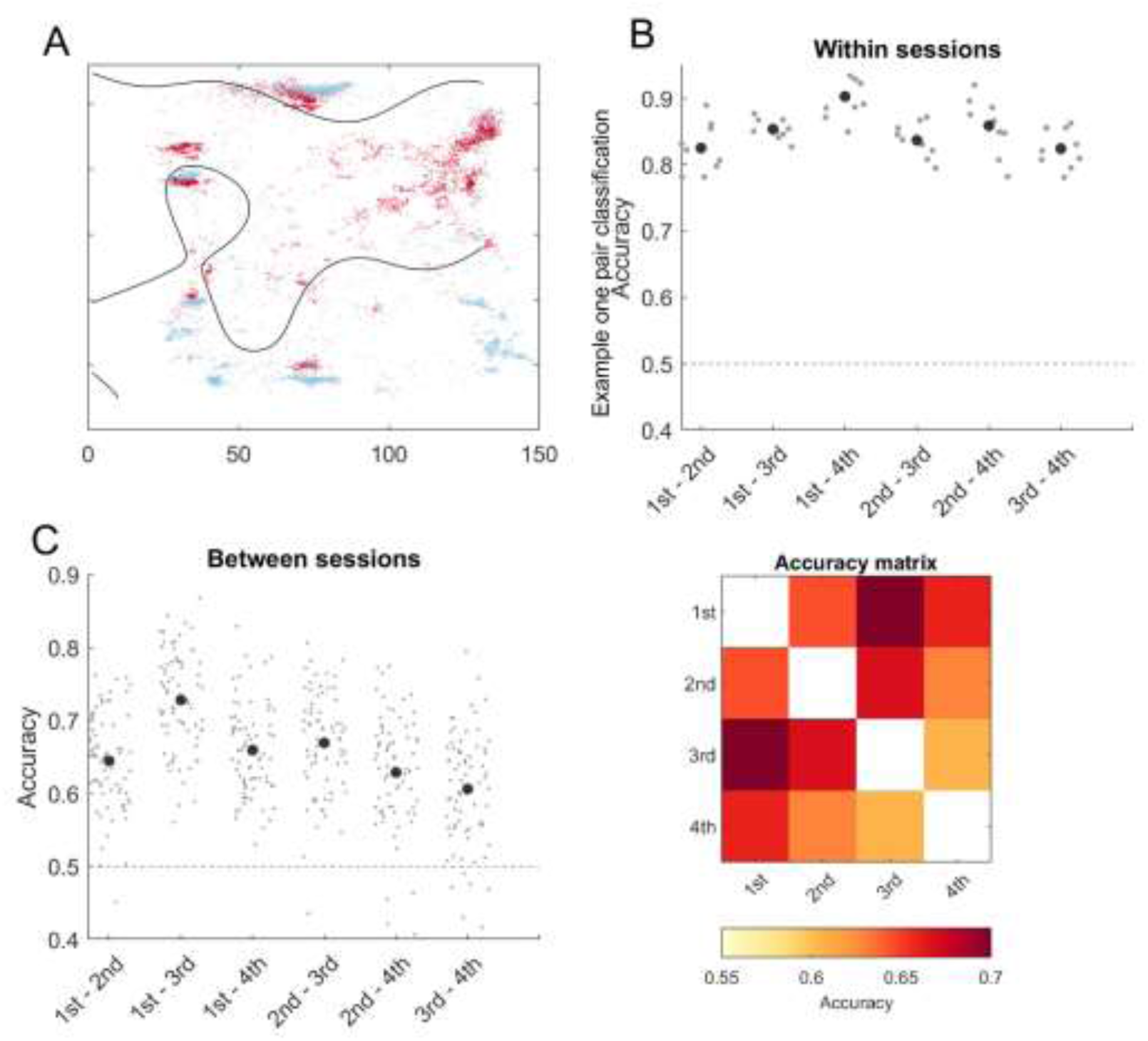
Decoding monkeys identity based on spatial occupancy in the second group of macaques. (A) Example of one such trained classifier. The trained classifier was tested in a cross validated manner or on all remaining 18 recording sessions. (B) To decode spatial occupancy within each session and monkey pair cross-validation procedure was performed. Decoding accuracy was above the chance level for all monkey pairs (bottom left panel), showing that each animal sustained its spatial footprint within each session. (C) The same decoding was performed on data across sessions which showed similar pattern of results. Decoding accuracy was above the chance level for all monkey pairs, demonstrating that each animal sustained its spatial footprint across multiple days. Average accuracy scores for each monkey pair are summarized in matrix form in the bottom right panel.

**Supplementary Figure 3.**
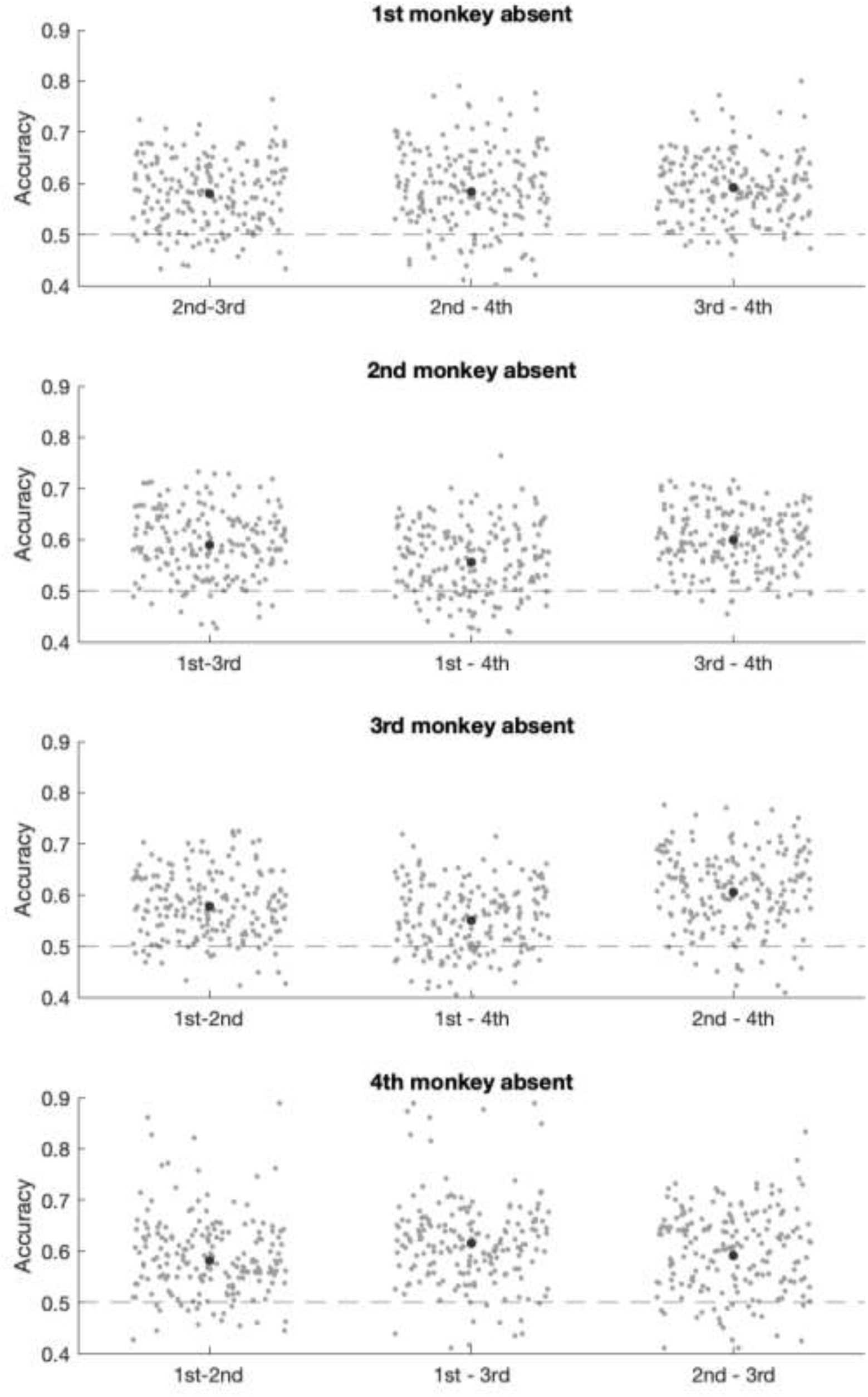
Decoding spatial occupancy between quadruple and triplet sessions. Classifiers were trained on quadruple sessions and tested on triplet sessions. Decoding accuracy was above the chance level for all monkey pairs, showing that each animal sustained its spatial footprint despite removal of one of the monkeys.

**Supplementary Figure 4.**
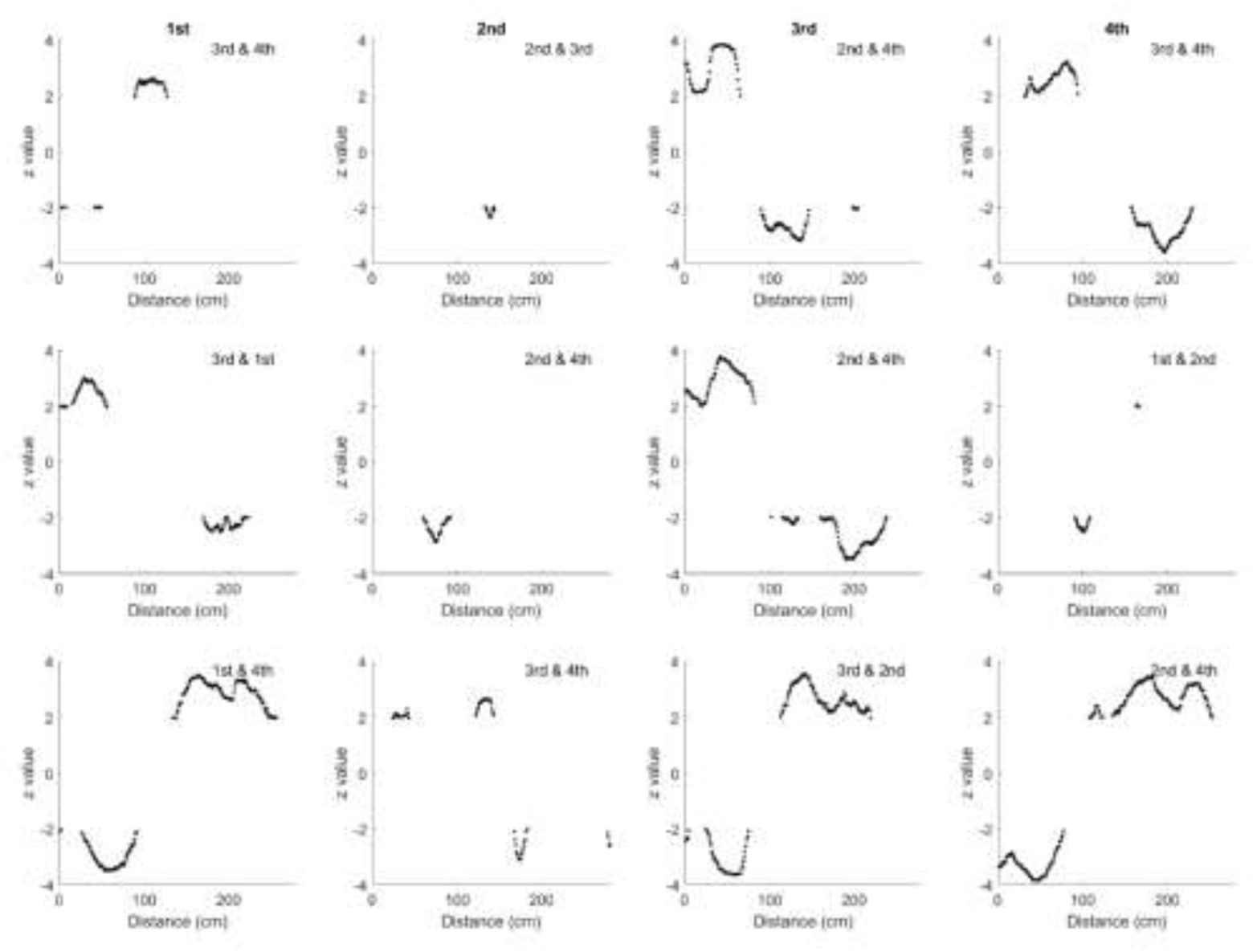
Comparison of density distributions plotted in Figure 4B. The columns of grid plot correspond to one monkey. The rows correspond to a second monkey from the pair. Time axis corresponds to time axis in Figure 4B.

**Supplementary Figure 5.**
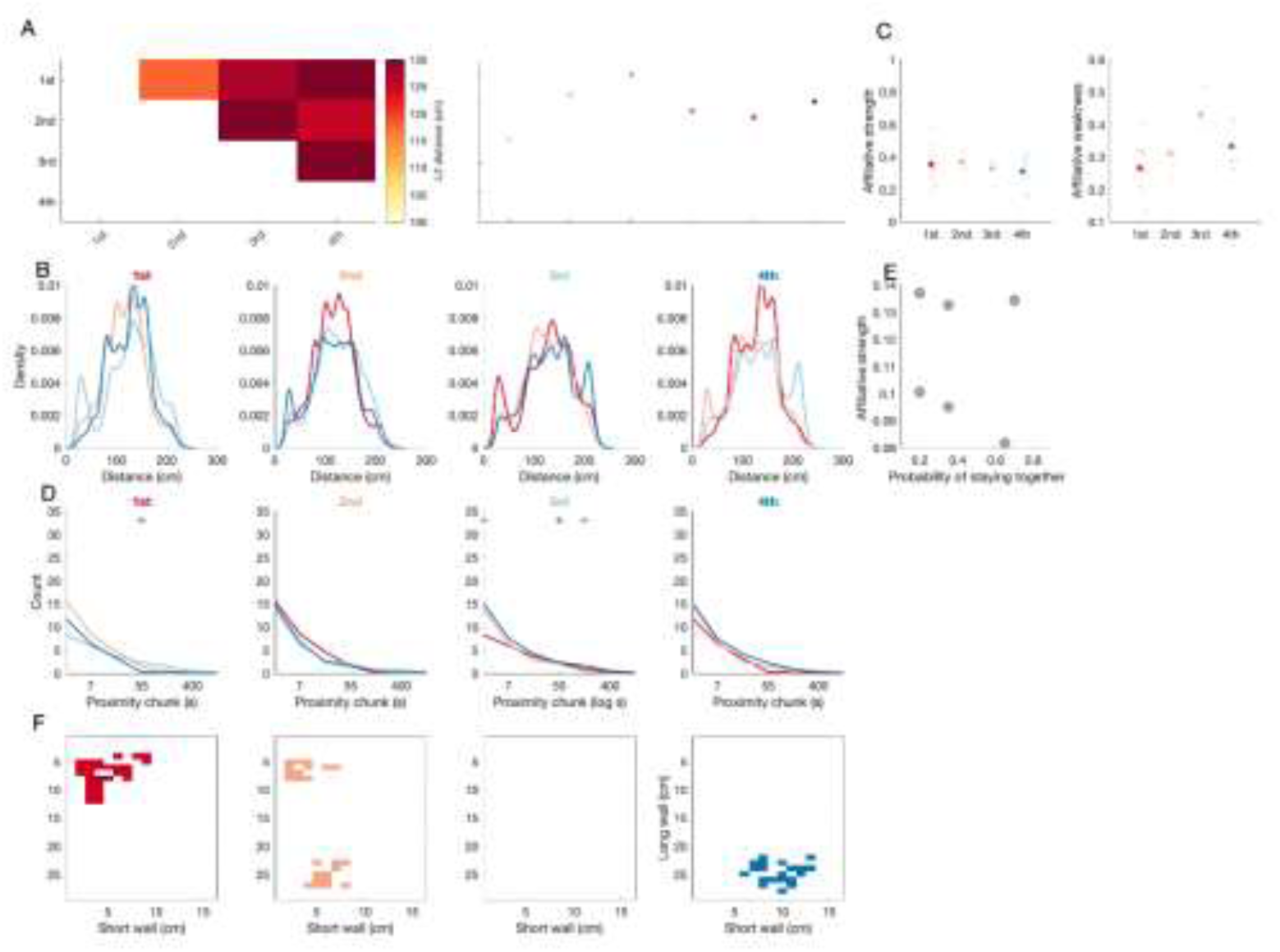
Features of interpersonal distance as a proxy of social structure in the second group of macaques. (A) The matrix represents the average distance for all sessions (n=9) for each pair of monkeys depicted in the matrix form. The redness of each cell represents average distance between a pair of monkeys. Higher values are obtained for 1^st^ and 4^th^ in the hierarchy (top right). (B) We computed Euclidian distance between each pair of monkeys over time. Individual density distribution depicts distribution of distance between given monkey pair. For example, the first panel displays three density functions which depict distribution of distance between 1st monkey and other three monkeys. The same follows for the three other plots. (C) Affiliative strength and affiliative weakness calculated as total time each monkey spent with all other monkeys or total time spend away from other monkeys, respectively. Monkeys did not differ in the values of affiliative strength. Affiliative weakness was lower for 1^st^ monkey as compared to all 3^rd^ and 4^th^ (3^rd^: Z = 2.83, *p* = 0.005; 4^th^: Z = 2.12, *p* = 0.034). The 2^nd^ monkey had lower affiliative weakness than 3^rd^ (4^th^: Z = 2.47, *p* = 0.013) but not 4^th^ (Z = 0.97, p > 0.05). 4^th^ monkey showed lower affiliative weakness than 3^rd^ monkey (Z = 2.30, *p* = 0.027). (D) Temporal distribution of close proximity chunks. The same color coding follows as in the panel A. The top stars indicate the samples where significant differences have been obtained. (E) Map of locations where monkeys was separated from the other monkeys by at least 120 cm.

**Supplementary Figure 6.**
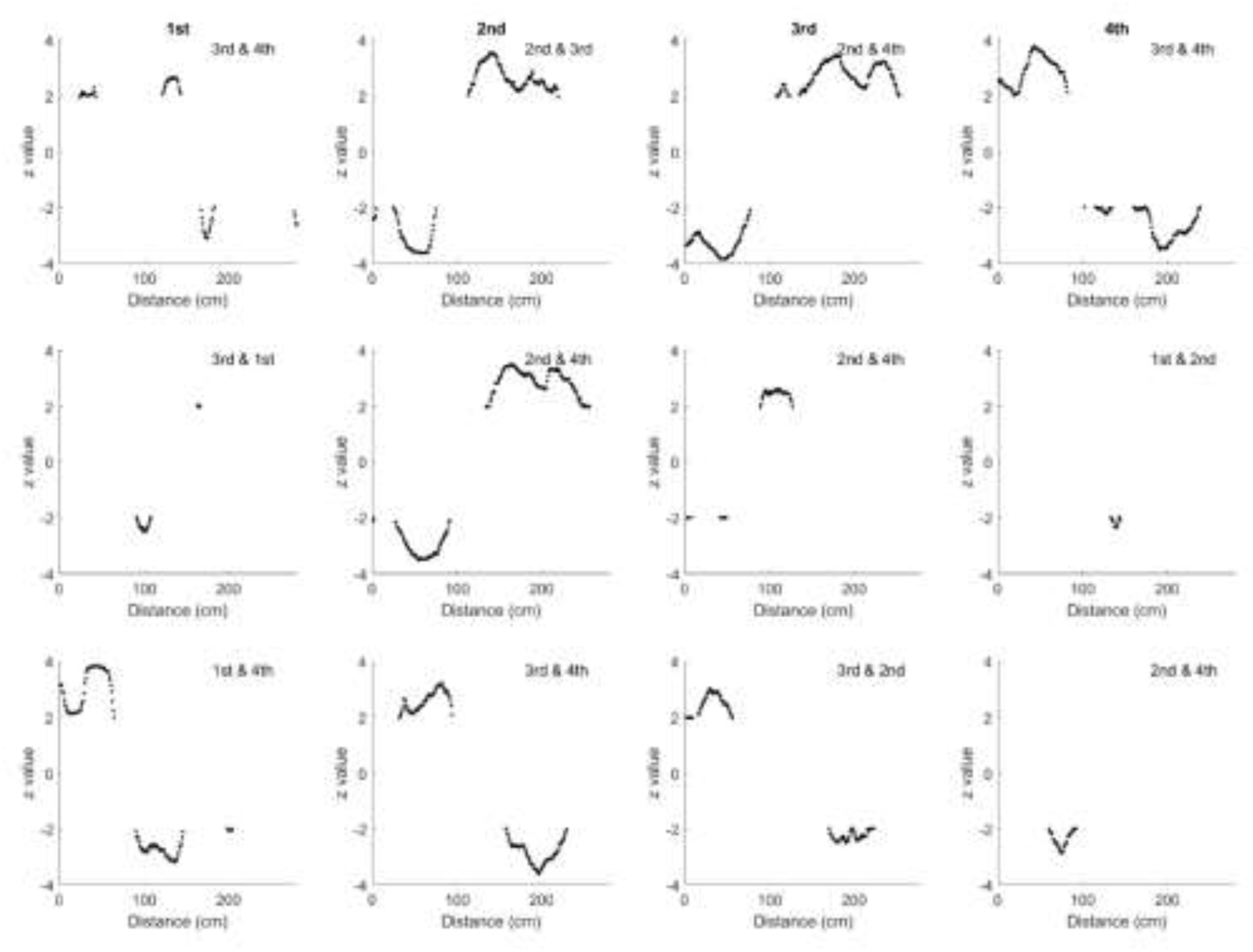
Comparison of density distributions plotted in Supplementary Figure 5B. The columns of grid plot correspond to one monkey. The rows correspond to a second monkey from the pair. Time axis corresponds to time axis in Supplementary Figure 5B and Figure 4A.

**Supplementary Figure 7.**
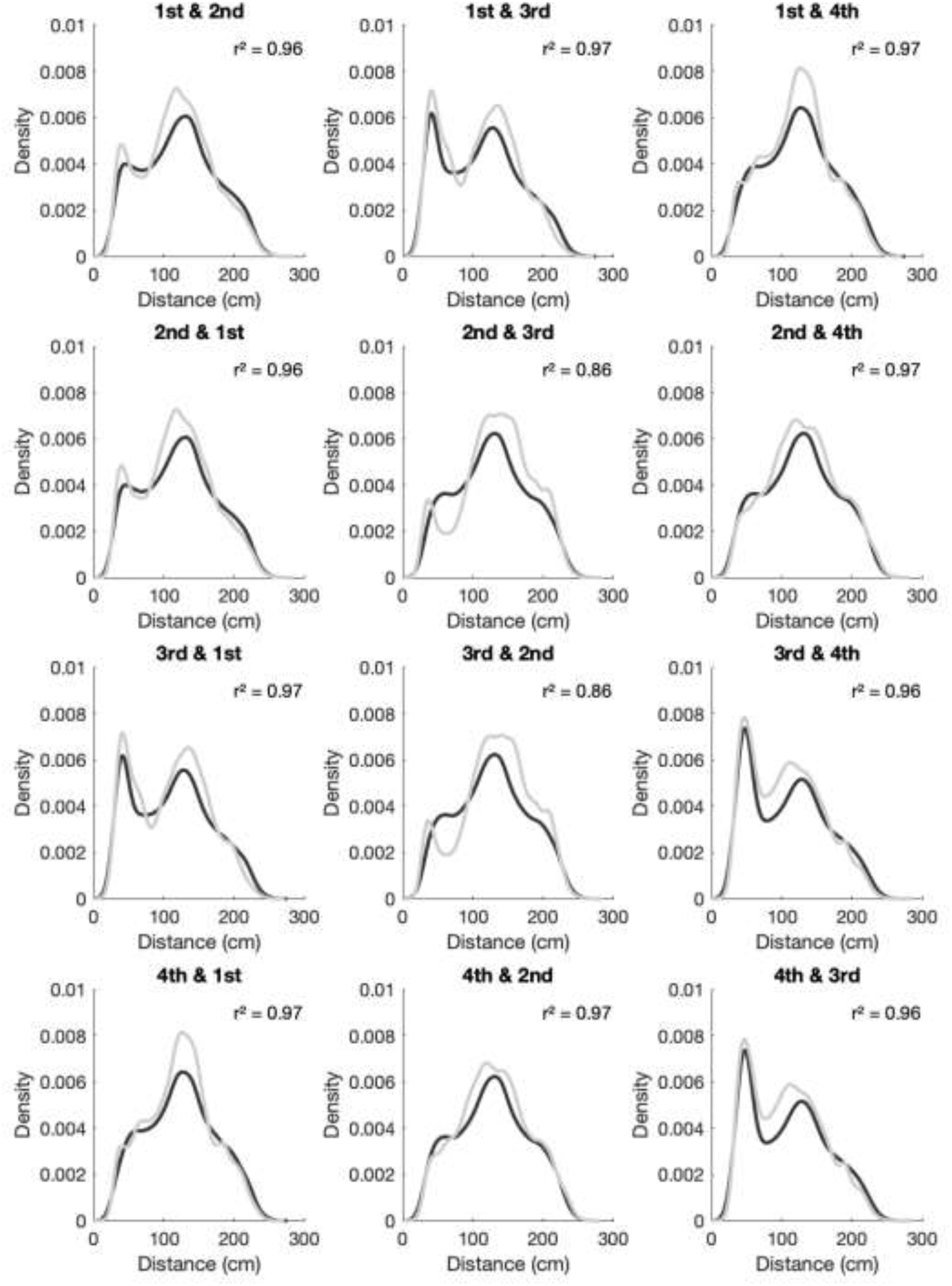
Model fits. Each distance distribution for a given monkey pair was fitted with simulated distribution with best fitting parameters. Empirical distributions are plotted in grey, whereas the simulated once in black for each pair of monkeys.

**Supplementary Figure 8.**
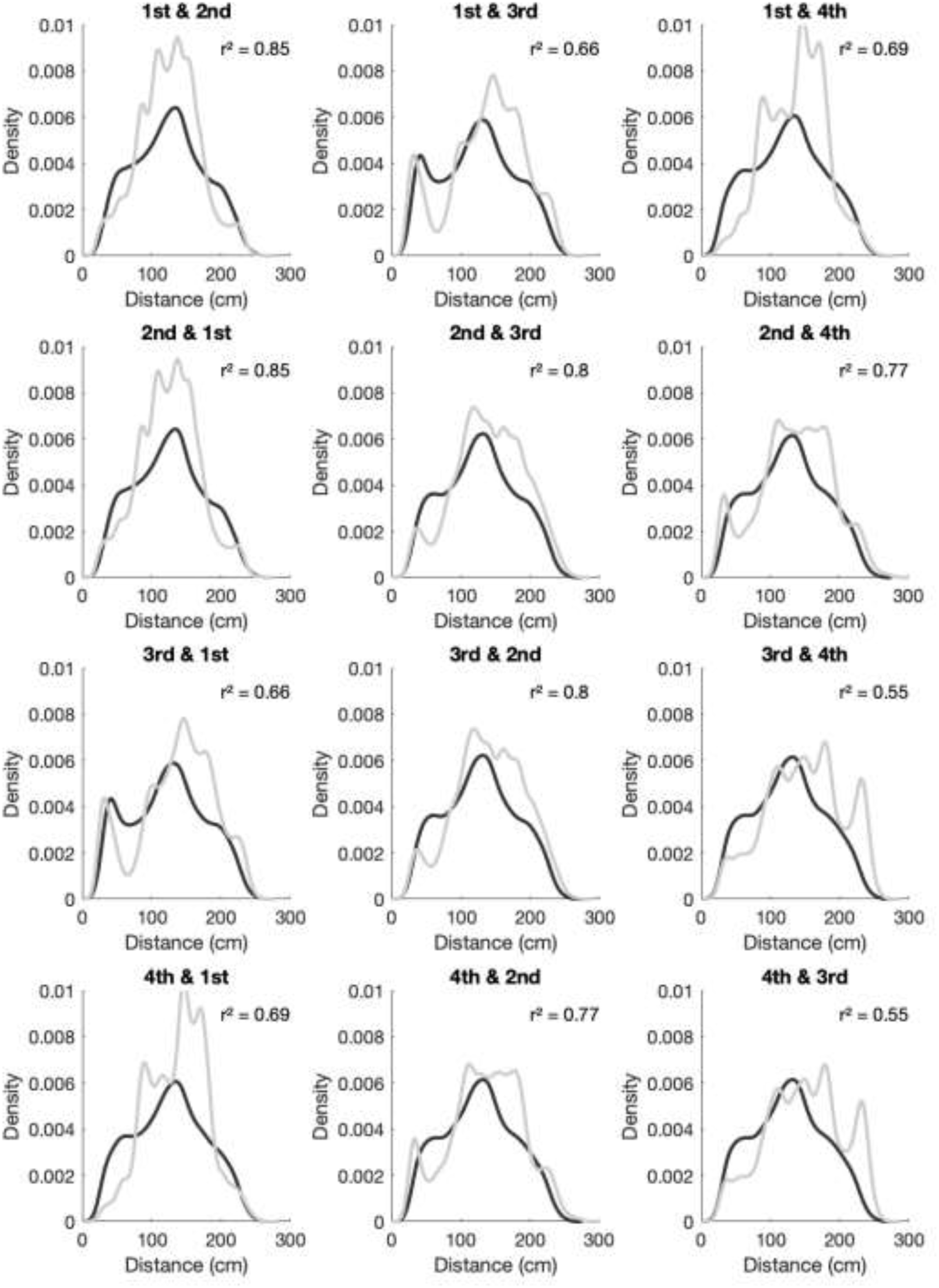
Model fits in the group of male macaques. All details remained the same as in the group of females.

## Methods

### Data collection

We recorded monkey positions within confined space (**Fig. 1AB**). We used two unisex groups of four female monkeys (*Macaca mulatta, age 4 to 5*). Monkeys where continuously recorded using 6 cameras. Each animal’s position was concurrently tracked with a sub-second resolution using collar’s color as an identifier (S. Ballesta et al. 2014). We replicated the results in the second group of four monkeys recorded in the same environment (*Macaca fascicularis, age 4 to 5*).

Both groups of 4 monkeys were housed together. For the first group of females (*M. mulatta*) we recorded data on 19 separate days. Each recording session took 3 hours. All the analyses were replicated in a second group of 4 animals (quadruple) for a total of 9 sessions.

To investigate the effects of social context on occupancy we recorded 10 sessions where only three monkeys reside inside the cage (triplet). As there were 4 monkeys that resulted in 4 kinds of triplet sessions.

### Assessment of dominance hierarchy

In order to characterize the dominance relationship individuals in the group, we performed 8 sessions of a food-grab test through which a food was offered to isolated pairs. this was the only way to assess beyond hierarchy for number 3 and 4, as when all four monkeys were together, only number 1 and 2 retrieved food. In our tests, the monkey which retrieved the food over all other 3 was termed #1, while the 2^nd^ retrieved the food over others only in the absence of the highest ranked monkey and so on. The hierarchy of group B was assessed using the “water bottle test” which scores hierarchy through a similar competition outcome method (Varley and Symmes 1966; Sébastien Ballesta et al. 2014).

### Decoding analyses

All decoding based analyses were performed using Support Vector Machine classifier (SVM) on the simultaneously recorded monkey positions using 3 dimensions, using *fitcsvm* MatLab function. We used radial basis function kernel. The ‘box constraint’ (aka ‘C parameter’) was set to 1. For within class (quadruples) we used 10-fold cross validation. For decoding assessing generalization between quadruple and triplet sessions we used entire session for the training and another session for testing.

### Computing social influence metrics

For each monkey, session, and condition we computed normalized occupancy maps expressing the proportion of time spent in a given part of the area. To summarize between session change we computed a single number index as L1 normed difference between normalized occupancy maps, depicted as single data point in **Fig. 2A**. We computed L1 normed difference between all combinations of quadruple and triplet sessions (**Fig. 2A**, colored column in each panel). Such index (*D*) shows amount of change between the quadruple and triplet context for a particular monkey, quantifying the amount of change for a given monkey under a particular social context. We also computed L1 normed difference between all combinations of quadruple sessions (**Fig. 2A**, black column in each panel), which we treat as a reference level for changes observed in triplet sessions.

As number of triplet sessions differed between monkeys, and thus also between context conditions, we accounted for that fact using a variant of permutation testing. For each permutation sample we randomly choose 100 quadruple and 100 triplet values of *D.* We repeat

that procedure 200 times, which resulted in a distribution of *t* and *p* values. We based our inference on average value of *p*.

### Assessment of affiliative strength

We first computed Euclidian distance for each sample for a given pair of monkeys, resulting in one time series for a given pair of monkeys (**Fig. 4A**). As such that analysis neglects the absolute spatial positioning of individuals. To compute the affiliative strength for each session we calculated the cumulative sum of distance for each pair of monkeys. We summed up this metric for all three pairs for each individual. We calculated cumulative some over distance ranging from 0 to 80 cm, which approximately captured the peak of close proximity. We then compared this metric using sessions as samples.

### Assessment of close proximity chunks

To identify the chunks of close proximity, where two individuals where separated by less than 50 cm we counted the consecutive number of samples that fall in that criteria. In case of an individual sample it was considered as a one sample chunk. To facilitate the visualization of results we convert the time scale to logarithmic scale and histograms where calculated. Finally, different histograms (**Fig. 4C**) where compared within different intervals using Kolmogorov-Smirnov test.

### Random walk model simulation

To simulate monkeys movement with random walks we set up 240 by 140 steps big lattice. Based on the empirical data we knew that monkeys spend most of their time on the boundaries of the colony room, thanks to usage o perches. Thus we constrained the lattice such that agents can move in the vicinity of 40 units, perpendicular the wall. At each step the agents would move in a random direction. To account for variable speed of movement the size of the step was randomized from uniform distribution from 1 to 40 in steps of 1. We simulated 10^4^ random walk samples. We than calculated Euclidian distance between a pair of agents.

In the second simulation we aimed at modeling the peak of close proximity as random walk model did not capture bimodal distributions. Therefore, we enhanced the random walk model with addition of a simple rule. On every sample that two agents were in close proximity (<10) the new parameter expressed the probability of staying in close proximity on the next sample (P(*StayTogether*)). We run separate simulations with the values of P(*StayTogether*) ranging from 0.1 to 0.9 in steps of 0.05.

### Calculation of private space

To find out distribution of private space for every given monkey we selected the data points when a given monkey was simultaneously separated from all the other monkeys for at least 150 cm. These selected data points were broken into a 29 by 16 matrix. Normalized maps where computed for each and every session. Finally, for each monkey a difference between her occupancy and averaged occupancy of other monkeys was computed, resulting in map specific for a given monkey. The unthresholded p-values where clustered using density based spatial clustering with *epsilon* and *minpts* parameters of 3 and 12. Thus the resulting map shows spatial distribution of private space where a given monkey tends to reside on her own.

## Bibliography

1. Ballesta, S., G. Reymond, M. Pozzobon, and J.-R. Duhamel. 2014. ‘A Real-Time 3D Video Tracking System for Monitoring Primate Groups’. Journal of Neuroscience Methods 234 (August):147–52. 10.1016/j.jneumeth.2014.05.022.

2. Ballesta, Sébastien, Gilles Reymond, Mathieu Pozzobon, and Jean-René Duhamel. 2014. ‘Compete to Play: Trade-Off with Social Contact in Long-Tailed Macaques (Macaca Fascicularis)’. Edited by Giuseppe Di Pellegrino. *PLoS ONE* 9 (12): e115965. 10.1371/journal.pone.0115965.

3. Bartumeus, Frederic, M. G E. Da Luz, G. M. Viswanathan, and J. Catalan. 2005. ‘ANIMAL SEARCH STRATEGIES: A QUANTITATIVE RANDOM-WALK ANALYSIS’.

4. Ecology 86 (11): 3078–87. 10.1890/04-1806.

5. Bates, Brian C. 1970. ‘Territorial Behavior in Primates: A Review of Recent Field Studies’.

6. Primates 11 (3): 271–84. 10.1007/BF01793893.

7. Bovet, Pierre, and Simon Benhamou. 1988. ‘Spatial Analysis of Animals’ Movements Using a Correlated Random Walk Model’. Journal of Theoretical Biology 131 (4): 419–33. 10.1016/S0022-5193(88)80038-9.

8. Burt, William Henry. 1943. ‘Territoriality and Home Range Concepts as Applied to Mammals’. Journal of Mammalogy 24 (3): 346. 10.2307/1374834.

9. Caselli, Christini B., Paulo H.B. Ayres, Shalana C.N. Castro, Antonio Souto, Nicola Schiel, and Cory T. Miller. 2018. ‘The Role of Extragroup Encounters in a Neotropical, Cooperative Breeding Primate, the Common Marmoset: A Field Playback Experiment’. Animal Behaviour 136 (February):137–46. 10.1016/j.anbehav.2017.12.009.

10. Chan, Stephanie, Hsieh Fushing, Brianne A. Beisner, and Brenda McCowan. 2013. ‘Joint Modeling of Multiple Social Networks to Elucidate Primate Social Dynamics: I. Maximum Entropy Principle and Network-Based Interactions’. PloS One 8 (2): e51903. 10.1371/journal.pone.0051903.

11. Codling, Edward A, Michael J Plank, and Simon Benhamou. 2008. ‘Random Walk Models in Biology’. Journal of The Royal Society Interface 5 (25): 813–34. 10.1098/rsif.2008.0014.

12. Dunbar, R.I.M. 1991. ‘Functional Significance of Social Grooming in Primates’. Folia Primatologica 57 (3): 121–31. 10.1159/000156574.

13. Farine, D. R., A. Strandburg-Peshkin, I. D. Couzin, T. Y. Berger-Wolf, and M. C. Crofoot. 2017. ‘Individual Variation in Local Interaction Rules Can Explain Emergent Patterns of Spatial Organization in Wild Baboons’. Proceedings of the Royal Society B: Biological Sciences 284 (1853): 20162243. 10.1098/rspb.2016.2243.

14. Feister, A.2018. ‘Nonhuman Primate Evaluation and Analysis Part 1: Analysis of Future Demand and supplyPrimate Evaluation and Analysis Part 1: Analysis of Future Demand and Supply’. Https://Orip.Nih.Gov/Nonhuman-Primate-Evaluation-and-Analysis-Part-1-Analysis-Future-Demand-and-Supply, 2018.

15. French, Jeffrey A., Colleen M. Schaffner, Rebecca E. Shepherd, and Marnie E. Miller. 1995. ‘Familiarity with Intruders Modulates Agonism towards Outgroup Conspecifics in Wied’s Black-tufted-ear Marmoset ( Callithrix Kuhli : Primates, Callitrichidae)’. Ethology 99 (1–2): 24–38. 10.1111/j.1439-0310.1995.tb01086.x.

16. Hansen, Malene F., Signe Ellegaard, Maria M. Moeller, Floris M. Van Beest, Agustin Fuentes, Ventie A. Nawangsari, Carsten Groendahl, et al. 2020. ‘Comparative Home Range Size and Habitat Selection in Provisioned and Non-Provisioned Long-Tailed Macaques (Macaca Fascicularis) in Baluran National Park, East Java, Indonesia’. Contributions to Zoology 89 (4): 393–411. 10.1163/18759866-bja10006.

17. Hayduk, Leslie A. 1978. ‘Personal Space: An Evaluative and Orienting Overview.’ Psychological Bulletin 85 (1): 117–34. 10.1037/0033-2909.85.1.117.

19. Hinsch, Martin, and Jan Komdeur. 2017. ‘What Do Territory Owners Defend Against?’ Proceedings of the Royal Society B: Biological Sciences 284 (1849): 20162356. 10.1098/rspb.2016.2356.

20. José-Domínguez, Juan Manuel, Marie-Claude Huynen, Carmen J. García, Aurélie Albert- Daviaud, Tommaso Savini, and Norberto Asensio. 2015. ‘Non-Territorial Macaques Can Range Like Territorial Gibbons When Partially Provisioned With Food’. Biotropica 47 (6): 733–44. 10.1111/btp.12256.

21. Lazaro-Perea, Cristina. 2001. ‘Intergroup Interactions in Wild Common Marmosets, Callithrix Jacchus: Territorial Defence and Assessment of Neighbours’. Animal Behaviour 62 (1): 11–21. 10.1006/anbe.2000.1726.

22. MacLean, Evan L., Sheila Roberts Prior, Michael L. Platt, and Elizabeth M. Brannon. 2009. ‘Primate Location Preference in a Double-Tier Cage: The Effects of Illumination and Cage Height’. Journal of Applied Animal Welfare Science 12 (1): 73–81. 10.1080/10888700802536822.

23. Maestripieri, Dario. 2007. Macachiavellian Intelligence: How Rhesus Macaques and Humans Have Conquered the World. University of Chicago Press. 10.7208/chicago/9780226501215.001.0001.

24. Maestripieri, Dario, and Christy L. Hoffman. 2012. ‘Behavior and Social Dynamics of Rhesus Macaques on Cayo Santiago’. In Bones, Genetics, and Behavior of Rhesus Macaques, edited by Qian Wang, 247–62. New York, NY: Springer New York. 10.1007/978-1-4614-1046-1_12.

25. Martel, Frances L., Claire M. Nevison, Michael J. A. Simpson, and Eric B. Keverne. 1995. ‘Effects of Opioid Receptor Blockade on the Social Behavior of Rhesus Monkeys Living in Large Family Groups’. Developmental Psychobiology 28 (2): 71–84. 10.1002/dev.420280202.

26. Matheson, Megan D., and Irwin S. Bernstein. 2000. ‘Grooming, Social Bonding, and Agonistic Aiding in Rhesus Monkeys’. American Journal of Primatology 51 (3): 177–86. 10.1002/1098-2345(200007)51:3<177::AID-AJP2>3.0.CO;2-K.

27. McCowan, Brenda, Brianne Beisner, and Darcy Hannibal. 2018. ‘Social Management of Laboratory Rhesus Macaques Housed in Large Groups Using a Network Approach: A Review’. Behavioural Processes 156 (November):77–82. 10.1016/j.beproc.2017.11.014.

28. Mertl-Millhollen, Anne S. 1988. ‘Olfactory Demarcation of Territorial but Not Home Range Boundaries by Lemur Catta’. Folia Primatologica 50 (3–4): 175–87. 10.1159/000156344.

29. Mitani, John C., David P. Watts, and Sylvia J. Amsler. 2010. ‘Lethal Intergroup Aggression Leads to Territorial Expansion in Wild Chimpanzees’. Current Biology 20 (12): R507–8. 10.1016/j.cub.2010.04.021.

30. Rebout, Nancy, Christine Desportes, and Bernard Thierry. 2017. ‘Resource Partitioning in Tolerant and Intolerant Macaques’. Aggressive Behavior 43 (5): 513–20. 10.1002/ab.21709.

31. Schino, Gabriele. 2001. ‘Grooming, Competition and Social Rank among Female Primates: A Meta-Analysis’. Animal Behaviour 62 (2): 265–71. 10.1006/anbe.2001.1750.

32. Seyfarth, Robert M. 1977. ‘A Model of Social Grooming among Adult Female Monkeys’. Journal of Theoretical Biology 65 (4): 671–98. 10.1016/0022-5193(77)90015-7.

34. Journal of Theoretical Biology. 1980. ‘The Distribution of Grooming and Related Behaviours among Adult Female Vervet Monkeys’. *Animal Behaviour* 28 (3): 798–813. 10.1016/S0003-3472(80)80140-0.

35. Simons, Noah D., Vasiliki Michopoulos, Mark Wilson, Luis B. Barreiro, and Jenny Tung. 2022. ‘Agonism and Grooming Behaviour Explain Social Status Effects on Physiology and Gene Regulation in Rhesus Macaques’. Philosophical Transactions of the Royal Society B: Biological Sciences 377 (1845): 20210132. 10.1098/rstb.2021.0132.

36. Thierry, Bernard. 2007. ‘Unity in Diversity: Lessons from Macaque Societies’. *Evolutionary Anthropology: Issues*, News, and Reviews 16 (6): 224–38. 10.1002/evan.20147.

37. Thierry, Bernard, M Singh, and W Kaumann. 2004. Macaque Societies: A Model for the Study of Social Organization. Cambridge University Press: Cambridge, MA. Vol. 41.

38. Varley, Margaret, and David Symmes. 1966. ‘The Hierarchy of Dominance in a Group of Macaques’. Behaviour 27 (1–2): 54–74. 10.1163/156853966X00100.

39. Willems, Erik P., Barbara Hellriegel, and Carel P. van Schaik. 2013. ‘The Collective Action Problem in Primate Territory Economics’. Proceedings of the Royal Society B: Biological Sciences 280 (1759): 20130081. 10.1098/rspb.2013.0081.

40. Xie, Pu-Zhen, Yu-Xuan Fan, Colin Chapman, Chi Ma, Cheng-Feng Wu, Ping Hu, Liu-Liu Hu, and Peng-Fei Fan. 2024. ‘Determinants of Macaques’ Space Use: A Test for the Ecological Constraints Model Using GPS Collars’. American Journal of Primatology 86 (8): e23636. 10.1002/ajp.23636.

